# Characterizing the genetic diversity of the Andean blueberry (*Vaccinium floribundum* Kunth.) across the Ecuadorian Highlands

**DOI:** 10.1101/2020.09.30.319681

**Authors:** Pamela Vega-Polo, Maria M. Cobo, Andrea Argudo, Bernardo Gutierrez, Jennifer Rowntree, Maria de Lourdes Torres

## Abstract

The Ecuadorian *páramo*, a high altitude tundra-like ecosystem, is a unique source of various ecosystem services and distinct biodiversity. Anthropogenic activities are associated with its fragmentation, which alters ecological factors and directly threatens resident species. *Vaccinium floribundum* Kunth., commonly known as Andean blueberry or *mortiño*, is a wild shrub endemic to the Andean region and highly valued in Ecuador for its berries, which are widely used in food preparations and hold an important cultural value. Since it is a wild species, *mortiño* could be vulnerable to environmental changes, resulting in a reduction of the size and distribution of its populations. To evaluate the extent of these effects on the *mortiño* populations, we assessed the genetic diversity and population structure of the species along the Ecuadorian highlands. We designed and developed a set of 30 homologous SSR markers and used 16 of these to characterize 100 *mortiño* individuals from 27 collection sites. Our results revealed a high degree of genetic diversity (H_E_=0.73) for the Ecuadorian *mortiño*, and a population structure analyses suggested the existence of distinct genetic clusters present in the northern, central and southern highlands. A fourth, clearly differentiated cluster was also found and included individuals from locations at higher elevations. We suggest that the population structure of the species could be explained by an isolation-by-distance model and can be associated to the geological history of the Andean region. Our results suggest that elevation could also be a key factor in the differentiation of *mortiño* populations. This study provides an extensive overview of the species across its distribution range in Ecuador, contributing to a better understanding of its conservation status. These results can assist the development of conservation programs for this valuable biological and cultural resource and for the *páramo* ecosystems as a whole.

## Introduction

*Vaccinium floribundum* Kunth., commonly known as *mortiño* or Andean blueberry, is a woody perennial shrub from the Ericaceae family. It is endemic to the Andean region in South America ranging from Venezuela to Bolivia and can be found between 1600 to 4500 masl [1]. *V. floribundum* grows in high-altitude ecosystems such as cool montane forests and *páramos* (tundra-like ecosystems) at temperatures ranging from 7 to 18 °C and displays adaptations to withstand frost and freezing conditions [1–3].

In Ecuador, *mortiño* is valued primarily for its black-purple fruit widely used for the elaboration of traditional drinks, ice creams, wines and preserves [1,4]. The fruits have high concentrations of bioactive compounds such as polyphenols, antioxidants, anthocyanins, and flavonoids with potential beneficial effects on human health. Phytochemical studies have reported anti-inflammatory, tumor suppressing and blood sugar regulation properties, and have been shown to inhibit lipidic accumulation in human cells [5–12].

*Mortiño* berries are hand-picked directly from the *páramos* and sold in local markets. To our knowledge, *V. floribundum* has not yet been domesticated and established as a crop. Despite various attempts, its propagation is problematic due to low rates of seed germination and a lack of knowledge regarding the factors required for its growth [13,14].

*V. floribundum* plays an important environmental and ecological role, being one of the first species that recover after bouts of deforestation and man-made fires in the *páramo* ecosystems [15]. This is partly due to its high regenerative capacity driven by propagation from roots and other woody structures [16]. However, the *páramo* is considered a fragile ecosystem overall due to low levels of primary productivity and slow natural succession, which makes any recovery after human intervention relatively slow [17,18]. Different anthropogenic activities have affected the *páramo* ecosystem during the past century, and its fragmentation in the Andean region has directly impacted the diversity of its species [19–23]. Studies have shown that habitat fragmentation processes are associated with the generation of patches that prevent seed and pollen dispersal between populations, resulting in the isolation and reduction of the populations’ size [24–29]. The effects of habitat fragmentation and other anthropogenic factors on the conservation status of *V. floribundum* in the Ecuadorian *páramo* are currently unknown.

The genetic diversity of a number of species of the genus *Vaccinium* has been extensively studied, (e.g. *V. corymbosum* and *V. macrocarpon*) with the use of RAPD, ISSR and SSR molecular markers, with a focus on plant breeding and horticultural crop programs [30–35]. However, genetic diversity studies of *Vaccinium* wild species, adapted to live in unique ecosystems like the *páramo*, are scarce. In a previous study, we used 11 heterologous (SSR) markers developed *for V. corymbosum* and found a moderate degree of *V. floribundum* genetic diversity in three provinces in northern Ecuador. However, this approach presented some limitations, mainly regarding the use of heterologous markers and the restricted sampling range [36]. Therefore, the aim of this study was to characterize the genetic diversity and population structure of *V. floribundum* with the use of homologous SSR markers across the complete distribution range of the species in Ecuador. For this purpose, we designed and developed 30 homologous SSR markers and used a subset of these (n=16) for the genetic characterization of 100 *mortiño* individuals from different localities across all ten provinces in the Ecuadorian Highlands. The data presented here contributes to our understanding of the genetic makeup of this species in Ecuador.

## Materials and methods

### Sampling and DNA extraction

From June 2017 to August 2018, a total of 100 samples from individual plants were obtained from 27 collection sites (CS) distributed across ten provinces in the Ecuadorian Highlands: Carchi, Imbabura, Pichincha, Cotopaxi, Bolivar, Tungurahua, Chimborazo, Cañar, Azuay, and Loja. Three to seven samples were collected at each site (S1 Table). Samples were collected between 2881 and 4160 masl and all individuals were georeferenced (geographical location and altitude) using a Garmin ETrex 10 Outdoor Handheld GPS Navigation Unit (Garmin International Inc.). For the purposes of this study, we considered the Ecuadorian Highlands latitudinal range (~657 km measured from north to south) and assigned each collection site to one of three regions: northern, central and southern (each region has a vertical extension of approximately 219 km). The northern region comprises individuals from CS1 (La Cofradia in Carchi) to CS11 (Sigchos in Cotopaxi), the central region includes individuals from CS12 (Quilotoa in Cotopaxi) to CS20 (Surimpalti in Cañar), and the southern region comprises individuals from CS21 (San Miguel in Cañar) to CS27 (Podocarpus in Loja) (Fig 1; S1 Table).

**Fig 1.**
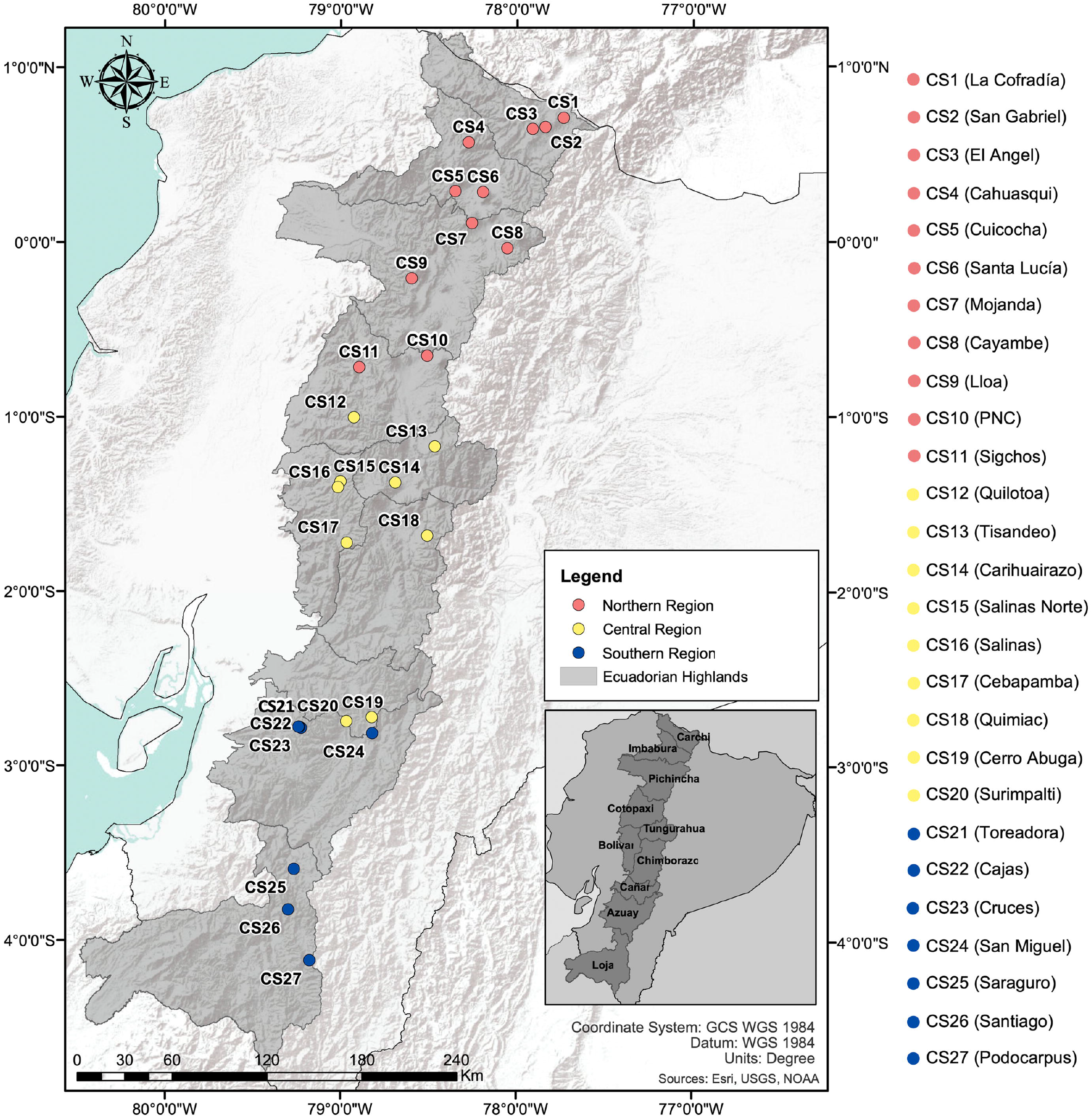
Map of Ecuador including the 27 *V. floribundum* collection sites (CS) in the Ecuadorian Highlands. *Mortiño* individuals sampled from 27 different collection sites (CS), distributed along three latitudinal regions (northern, central and southern) represented by different marker colors in the map. The sampling range included all ten provinces of the Ecuadorian Highlands showed in the bottom right.

Five to seven young leaves were collected from each individual and transported at 4°C to the Plant Biotechnology Laboratory at Universidad San Francisco de Quito, where they were stored at −20°C. Total genomic DNA was extracted from frozen leaves using the CTAB method [37]. DNA was quantified by spectrophotometry using a Nanodrop 2000 (Thermo Scientific).

### Development of SSR markers for *V. floribundum*

SSR markers were developed from a single DNA extraction of a *V. floribundum* individual using the Galaxy-based bioinformatics pipeline reported by Griffiths et al (2016) [38] at the University of Manchester genomics facility. The Illumina MiSeq platform was used with the shotgun 2 x 250 paired-end sequencing methodology (Nextera DNA Preparation Kit, Illumina, USA). The sample used 0.33 of a flow cell and primer design was optimized for use with Platinum Taq DNA polymerase (Invitrogen, USA) with an optimal melting temperature (T_m_) of 62°C (Min 59°C; Max 65°C) and a maximum difference among primer pairs of 3°C.

A total of 2 x 1,680,936 raw sequence reads was produced, with none flagged as poor quality. Sequence length ranged from 50 – 300 bp with a reported %GC content of 39. After screening, a total of 358 primer pairs were suggested, all amplifying SSRs with simple motifs. The majority of SSRs (333) were 2 bp motifs, but there were 22 with 3 bp motifs, and three with 4, 5 and 6 bp motifs, respectively.

From the initial 358 microsatellites markers, the first 30 were selected as candidates for our study; 25 with three or more base pairs motifs (3 – 6 bp), and five with 2 bp motifs. The melting temperature (Tm) of the primer sets ranged from 58°C to 63°C, with lengths between 18 and 25 bp (S2 Table). The primers were synthetized, including a universal tail (Tail A) in the locus-specific forward primers for genotyping. The universal tail A is complementary to fluorescently labelled universal primers (VIC, 6-FAM, NED or PET), which allows them to be incorporated during the amplification, obtaining a fluorescently labelled amplicon ready to be genotyped [39].

### Sample preparation and genotyping

Total genomic DNA (20 ng) was amplified in a 30μl reaction containing 1X PCR buffer, 2mM MgCl_2_, 0.15μM modified locus-specific forward primer, 0.5μM reverse primer, 0.5μM universal Tail A with a fluorophore (VIC, 6-FAM, NED or PET), 0.2mM dNTPs, and 1U Platinum Taq Polymerase. The Tail A primers label the amplified products at the 5’ end through a three-primer system as reported by Blacket et al (2012) [39], as the modified locus-specific forward primer is exhausted in early cycles and the fluorophore (VIC, 6-FAM, NED or PET) gets incorporated into the PCR fragments in the subsequent cycles.

Amplification consisted of 15 min at 94°C, followed by 40 cycles of 30 sec at 94°C, 90 sec at the standardized annealing temperature (58 – 63°C) (S2 Table), 60 sec a 72°C, and a final elongation of 5 min at 72°C. Successful PCR amplification was identified in 1.5% agarose gel electrophoresis, where one or two bands per locus were present. Labelled amplified products were genotyped by Macrogen (Seoul, Korea) on an ABI 3100 Genetic Analyzer (Applied Biosystems) automatic capillary sequencer, using 500 LIZ as a size standard.

### Genetic analyses

#### SSR marker performance, genetic diversity and population genetics

GeneMarker software (SoftGenetics LLC) was used to identify individual alleles and allele size. Polymorphic information content values for each SSR locus were obtained with the *polysat* R-based statistical package [40]. Expected heterozygosity (H_E_), observed heterozygosity (H_O_) and the number of alleles (N_A_) per locus and per collection site were estimated using the *adegenet* [41] and *hierfstat* [42] R packages. Private alleles were identified using the *poppr* R package [43], while null allele frequencies for all the SSR loci were determined with the FreeNA software through the EM algorithm [44]. Mean allelic richness (A_R_), standardized to de minimum sample size (n=3) through rarefaction, was estimated using the *diveRsity* R package [45]. Fixation indices (F_IS_) were calculated using the *adegenet* R package [41] and described as estimates of inbreeding.

#### Population structure and genetic differentiation

A principal coordinate analysis (PCoA) was performed with the *ade4* package [46] and the first three components plotted using the *ggplot* package [47]. In addition, a Bayesian analysis with an admixture model was performed using the program STRUCTURE 2.3.4 [48]. The potential number of genetic clusters (K) was evaluated between 1 and 10, with 10 independent repetitions for each K value. A 100,000 step burn-in period was used, followed by 1’000,000 Markov Chain Monte Carlo (MCMC) steps. The optimum value of K was evaluated through de Evanno method [49] using the web-based program STUCTURE HARVESTER [50]. The independent replicates for the optimum value of K were aligned with the program CLUMPP [51] and the final STRUCTURE graph was plotted with *DISTRUCT* [52].

The R package *poppr* was used to perform an analysis of molecular variance (AMOVA) to evaluate genetic differentiation among and within the identified genetic clusters [43]. Pairwise F_ST_ genetic distances between these clusters were estimated with the *hierfstat* package [42] using the Weir & Cockerham equation [53]. The same genetic distances, corrected for null alleles, were calculated with the FreeNA software [44]. Expected heterozygosity (H_E_), observed heterozygosity (H_O_), number of alleles (N_A_), number of private alleles (N_PA_) and the mean allelic richness (A_R_) standardized (n=14) through rarefaction, were also calculated for each of the clusters. The differences in the H_E_ between clusters were calculated through Monte-Carlo tests using *adegenet* R package [41]. Bonferroni corrections for multiple paired comparisons were applied. Furthermore, directional relative migration was calculated and plotted using the *diveRsity* R-based package [45], based on Nei genetic distances, to better describe the patterns of gene flow between the identified genetic clusters.

#### Relationship between elevation, geographic distances and genetic variability of *V. floribundum*

Given the geographic coordinates and the elevation (masl) of each sampled individual, a Mantel test was performed (10 000 permutations) using the *ape* R statistical package [54], to evaluate the relationship between the genetic (Nei’s) and geographic distances among the sampled individuals. Furthermore, to determine whether elevation was associated with the genetic diversity of *V. floribundum, stats* [55] and *ggpubr* R [56] statistical packages were used to perform a Pearson correlation test between the expected heterozygosity (H_E_) of individuals found at any given collection site and the mean elevation of said site.

## Results

### Validation of a set of homologous SSR markers developed for *V. floribundum*

Of the 30 SSR markers tested, four were excluded due to the absence of PCR products, and an additional 10 presented different artifacts like stutter peaks and excessive baseline noise. The remaining 16 homologous SSR markers showed reliable PCR amplification and highly fluorescent peaks (>1000 relative fluorescent units) after genotyping, and were selected for further analysis. These markers proved to be polymorphic and highly informative with an average polymorphic information content (PIC) value of 0.69. Mo025 (PIC=0.90) and Mo020 (PIC=0.88) were the most polymorphic markers, while Mo011 was the least polymorphic (PIC=0.26) (Table 1). A total of 179 alleles were identified (NA), with an average number of 11.2 alleles per locus (ranging from 6 alleles for Mo002 to 20 alleles for Mo020 and Mo025) (Table 1). Null alleles presented low to mid frequencies across all the SSR loci, ranging from 0.034 for Mo002 to 0.225 for Mo025 (S3 Table). Expected heterozygosity (H_E_) estimates ranged from 0.26 for Mo001 to 0.91 for Mo025 (Table 1).

**Table 1.** Genetic diversity indices of the 100 *V. floribundum* individuals analyzed with 16 SSR markers.

| Locus | T (°C) | Expected<br>size (bp) | $N_A$ | $H_O$ | $H_E$ | PIC |
| --- | --- | --- | --- | --- | --- | --- |
| Mo001 | 58 | 300-400 | 10 | 0.61 | 0.82 | 0.79 |
| Mo002 | 58 | 100-200 | 6 | 0.39 | 0.58 | 0.49 |
| Mo004 | 58 | 300-400 | 9 | 0.45 | 0.79 | 0.77 |
| Mo005 | 58 | 200-300 | 12 | 0.46 | 0.87 | 0.85 |
| **Mo007** | 63 | 200-300 | 9 | 0.35 | 0.58 | 0.50 |
| **Mo008** | 63 | 300-400 | 11 | 0.41 | 0.77 | 0.74 |
| **Mo009** | 63 | 400-500 | 7 | 0.25 | 0.59 | 0.55 |
| **Mo010** | 63 | 250-350 | 11 | 0.32 | 0.66 | 0.62 |
| **Mo011** | 58 | 300-400 | 6 | 0.22 | 0.26 | 0.26 |
| **Mo015** | 60 | 200-300 | 8 | 0.24 | 0.63 | 0.57 |
| **Mo016** | 60 | 350-450 | 13 | 0.45 | 0.79 | 0.76 |
| **Mo018** | 60 | 200-300 | 9 | 0.42 | 0.72 | 0.69 |
| **Mo020** | 60 | 300-400 | 20 | 0.59 | 0.89 | 0.88 |
| **Mo021** | 60 | 300-400 | 16 | 0.49 | 0.88 | 0.87 |
| **Mo024** | 60 | 300-400 | 12 | 0.56 | 0.88 | 0.87 |
| **Mo025** | 60 | 250-350 | 20 | 0.46 | 0.91 | 0.90 |
| **Mean** |  |  | **11.20** | **0.42** | **0.73** | **0.69** |

$N_A$ (number of alleles); $H_O$ (observed heterozygosity); $H_E$ (expected heterozygosity); and PIC (polymorphic information content). Optimum annealing temperature (in °C) for each set of primers and the expected size of the amplified band (or the allele found) are also presented. Primers sequences and SSR motifs are presented in the S2 Table.

### Genetic diversity of *V. floribundum* in the Ecuadorian Highlands

After genotyping 100 individuals from all the collection sites, an average number of 41.7 unique alleles per collection site were identified. CS18 (Quimiac) (N_A_=65; H_E_=0.6) and CS14 (Carihuairazo) (N_A_=60; H_E_=0.61), in the central region, presented the highest number of alleles and expected heterozygosity estimates. In contrast, CS24 (Cruces) (N_A_=28; H_E_=0.28) and CS23 (Cajas) (N_A_=26; H_E_=0.24) in the southern region, and CS12 (Quilotoa) (N_A_=26; H_E_=0.21) in the central region presented the lowest genetic diversity indicators (S4 Table). Similar results were found when analyzing the mean allelic richness (A_R_) per collection site, an estimate that corrects for the sampling imbalance between collection sites (S4 Table). The global expected heterozygosity (H_E_=0.73) (Table 1) revealed a high degree of genetic diversity for *V. floribundum* in the Ecuadorian Highlands.

### Population structure and genetic differentiation

A principal coordinate analysis (PCoA) was conducted to test genetic similarities between individuals and to define possible groupings. Two-dimensional plots were obtained (Fig 2) for the first three components explaining 42.83% of the total variance (PC1=21.25%; PC2=13.36%; PC3=8.22%). Overall, no clear differentiation between regions was observed; nonetheless, most of the individuals sampled in the northern region separated from the individuals sampled in the southern region through the variance explained by the second component (PC2) (Fig 2). Individuals from the central region were distributed along the PC2. Furthermore, a clearly differentiated group was identified (marked by a green ellipse, Fig 2) which included individuals from CS12 (Quilotoa) in the central region, and CS22, CS23, and CS24 (collection sites in Azuay) in the southern region. Individuals from this group were mainly separated from the others through the variance explained by the first component (PC1) (Fig 2).

**Fig 2.**
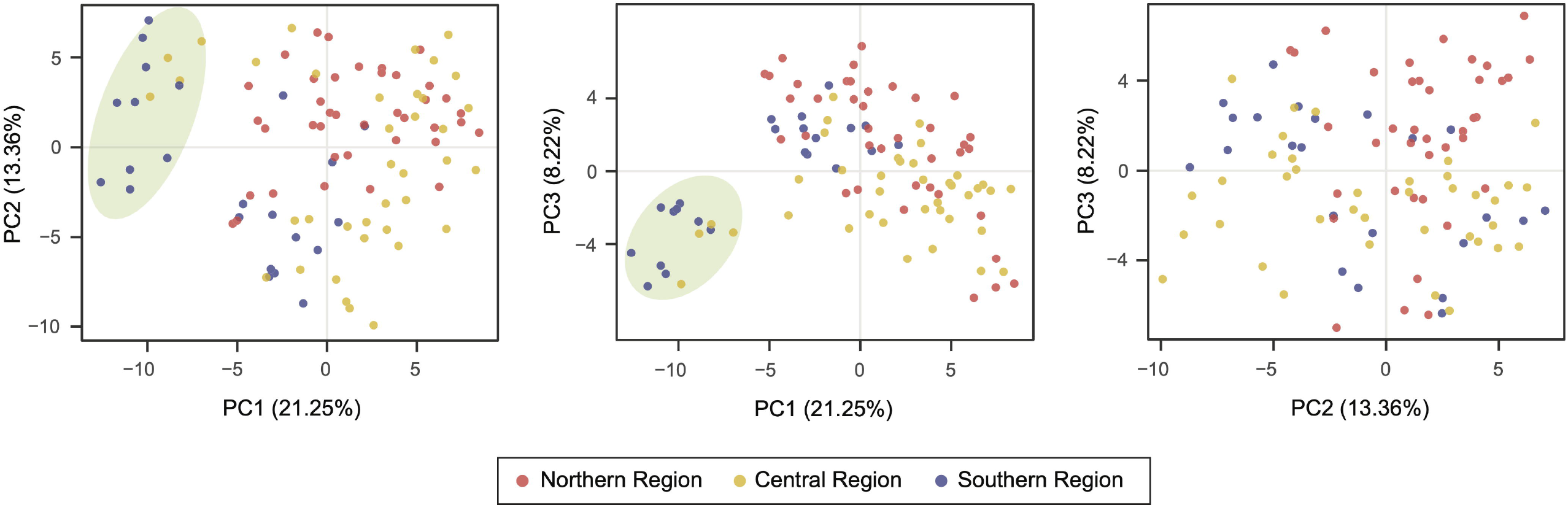
Principal coordinate analysis (PCoA) of V. *floribundum* using 16 SSR markers. The first three components represent 42.83% of the total variation. Individuals were assigned to their defined latitudinal region (northern, central and southern). One main group was identified (green ellipse) and was separated by the first component (PC1).

A Bayesian analysis was performed to better evaluate the genetic structure of *V. floribundum* in the Ecuadorian Highlands. The STRUCTURE results showed four possible genetic lineages (*K*=4), based on the highest Δ*K* calculated through the Evanno method [49] (S5 Table). Individuals were assigned to four clusters based on their predominant inferred ancestry (Fig 3A). Cluster 1 contained most of the individuals from the northern region (in red), cluster 2 included the individuals from the central region plus CS9 (Lloa) and CS11 (Sigchos) from the northern region (in yellow), and cluster 3 comprised the individuals from the southern region plus CS19 (Cerro Abuga) and CS20 (Surimpalti) from the central region (in blue) (Fig 3A). Notably, cluster 4 was highly consistent with the group identified in the PCoA (Fig 2), comprising the individuals from CS12 (Quilotoa in Cotopaxi) in the central region, and CS22, CS23 and CS24 (collection sites in Azuay) in the southern region (in green) (Fig 3A). When the number of possible genetic lineages was reduced to *K*=3 (Fig 3B) and *K*=2 (Fig 3C), cluster 4 remain grouped separately, showing a clear differentiation of this cluster from the others.

**Fig 3.**
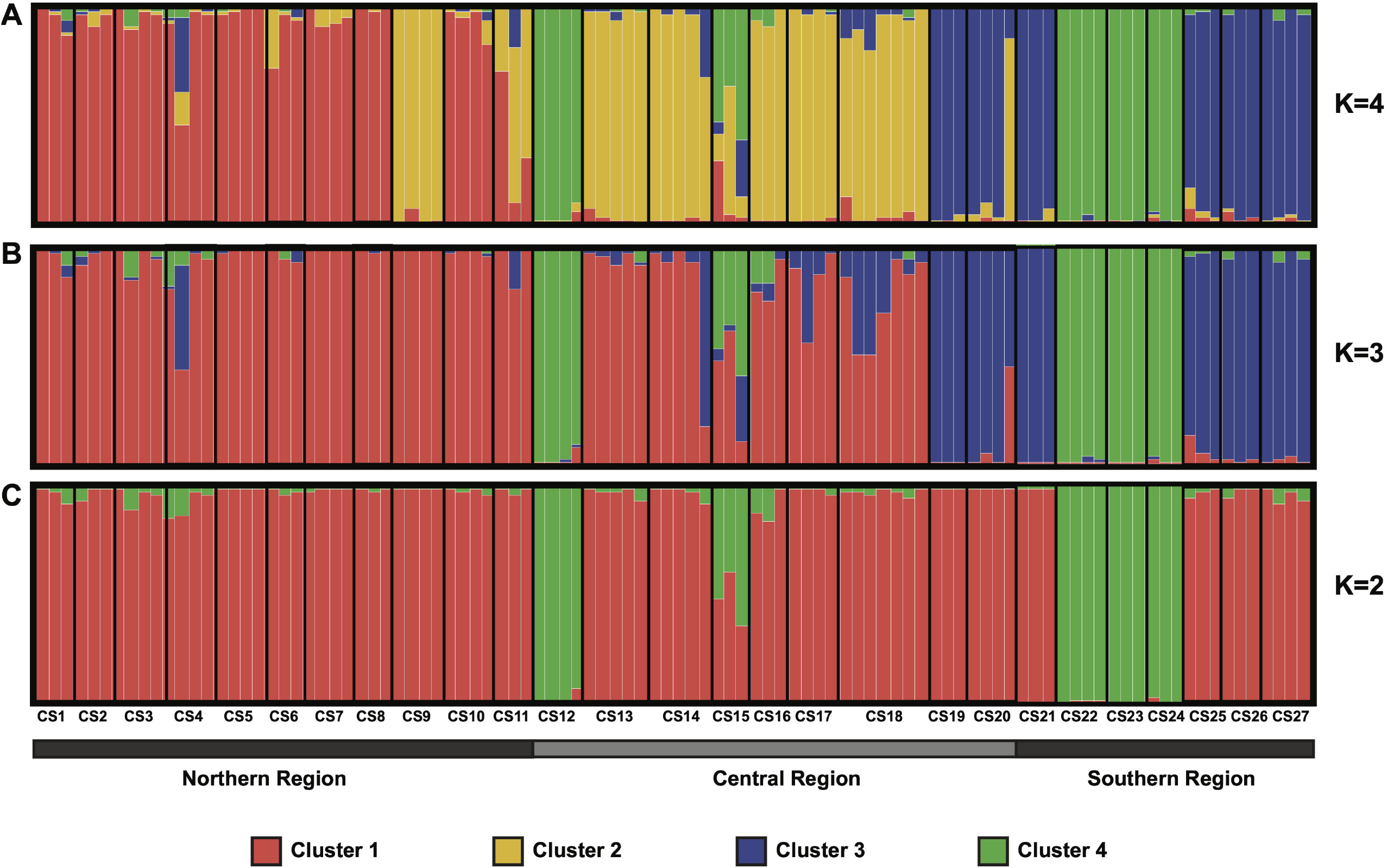
Bayesian analysis of the population structure of *V. floribundum* genotyped at 16 SSR loci under the Admixture model. *K* is the number of genetic lineages represented by different colors. The individual assignment probabilities given the optimal number of lineages (*K*=4, based on Δ*K*= 110.9) (A) are compared to the assignment probabilities for *K*=3 (B) and for *K*=2 (C). Individuals on the plots are ordered from left to right based on their sampling locations along the northern to southern axis (of the Ecuadorian Highlands), from CS1 (La Cofradia in Carchi), to CS27 (Podocarpus in Loja) (S1 Table).

An analysis of molecular variance (AMOVA) shows most of the variation (78.64%) occurs within the genetic clusters, and only 21.36% between these clusters (*p*=0.001) (S6 Table). Corrected F_ST_ genetic distances show the greater differentiation occurs between cluster 4 and clusters 1 (F_ST_=0.171), 2 (F_ST_=0.249) and 3 (F_ST_=0.230). The lowest genetic distance was found between clusters 1 and 2 (F_ST_=0.078), suggesting higher genetic similarities between these clusters (Table 2).

**Table 2.** Pairwise F_ST_ values between the four genetic clusters identified for *V. floribundum* in the Ecuadorian Highlands.

|  | Cluster 1 | Cluster 2 | Cluster 3 |
| --- | --- | --- | --- |
| Cluster 2 | 0.078 | - | - |
| Cluster 3 | 0.114 | 0.120 | - |
| Cluster 4 | 0.171 | 0.249 | 0.230 |

The $F_{ST}$ were estimated according to Weir and Cockerham (1984) and corrected for null alleles through FreeNA software. Correction for null alleles decreases the possibility of bias in the estimation of genetic differentiation values. Higher values indicate greater genetic differentiation between the analyzed clusters.

Finally, genetic diversity indices for each cluster were calculated (Table 3). The number of alleles (N_A_) ranged from 57 alleles for cluster 4 to 120 alleles for the cluster 2. The private alleles for each genetic cluster ranged between 8 (14.03%) for cluster 4 and 25 for the cluster 2 (20.83%). The highest expected heterozygosity (H_E_) estimate was observed for clusters 1 (H_E_=0.67), 2 (H_E_=0.65) and 3 (H_E_=0.68); with cluster 4 showing a significantly lower genetic diversity (H_E_=0.39, *p*=0.001) (Table 3). No statistical differences in H_E_ were found between cluster 1 and both clusters 2 (*p*=0.259) and 3 (*p*=0.777), nor between cluster 2 and cluster 3 (*p*=0.331). Among all individual collection sites, locations assigned to cluster 4 (CS12, CS22-CS24) presented the lowest diversity indices (S4 Table).

**Table 3.** Genetic diversity indices of the four *V. floribundum* identified genetic clusters.

| Clusters | N | N<sub>A</sub> | N<sub>PA</sub> | A<sub>R</sub> | H<sub>O</sub> | H<sub>E</sub> |
| --- | --- | --- | --- | --- | --- | --- |
| Cluster 1 | 35 | 118 | 18 | 2.25 | 0.45 | 0.67 |
| Cluster 2 | 31 | 120 | 25 | 2.09 | 0.48 | 0.65 |
| Cluster 3 | 20 | 103 | 15 | 2.03 | 0.51 | 0.68 |
| Cluster 4 | 14 | 57 | 8 | 1.65 | 0.08 | 0.39 |

N (number of individuals); N<sub>A</sub> (number of alleles); N<sub>PA</sub> (number of private alleles); A<sub>R</sub> (mean allelic richness calculated using rarefaction); H<sub>O</sub> (observed heterozygosity); and H<sub>E</sub> (expected heterozygosity) are given for all 100 individuals genotyped at 16 SSR loci distributed in four genetic clusters.

### Directional genetic differentiation and relative migration between *V. floribundum* genetic clusters

A directional relative migration network was inferred for the four genetic clusters, given their spatially structured distribution. These results revealed different levels of gene flow and possible migration rates between the four genetic clusters (Fig 4). As observed from genetic distances, clusters 1 and 2 presented the greatest genetic similarity (represented by the shorted distances between nodes). In contrast, cluster 4 presented the greatest differentiation in relation to the others, again in agreement with the results obtained by the genetic distances’ analyses (Table 2). A relatively high rate of potential bidirectional asymmetric gene flow was found between clusters 1 and 2, being higher from cluster 2 to cluster 1 than in the opposite direction. In turn, both clusters present moderate rates of bidirectional asymmetric gene flow with cluster 3. Remarkably, none of the clusters shows evidence of gene flow with cluster 4, which in turn presents minimal unidirectional gene flow with the other clusters (Fig 4).

**Fig 4.**
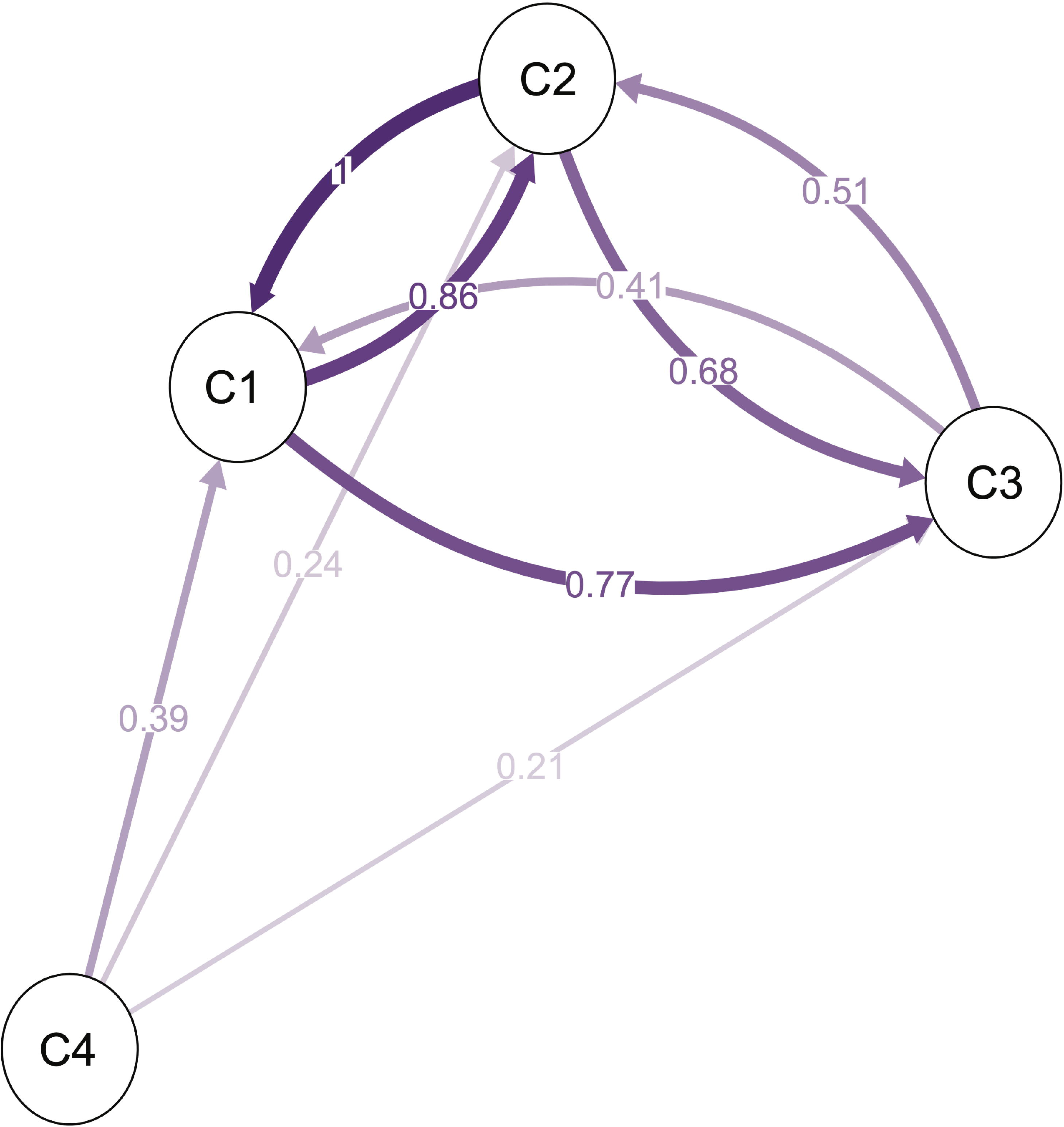
Directional relative migration network among the four genetic clusters found in the population structure analysis. The distances between nodes are proportional to the genetic similarity (Nei’s Gst) between populations, while the connecting lines shading reflect the relative migration rate between the populations. Numbers indicate the gene flow rate (scaled from 0 to 1).

### Relationship between elevation, geographic distances and genetic variability of *V. floribundum*

Mantel test results revealed a significant correlation between genetic and geographic distances (*r*=0.29; *p*=0.0001) between individuals, suggesting that a possible isolation by distance dynamic might explain the observed geographic structure of the studied populations. Furthermore, we observed a significant negative correlation between elevation and genetic diversity (evaluated through expected heterozygosity estimates) among the different collection sites through the three regions (*r*=-0.53; *p*=0.0045) (Fig 5; S7 Table). The regression equation (H_E_=0.968–0.00015*Elevation) can be approximated as a mean genetic diversity decrease of 0.00015 per every 1-meter increment in elevation along our altitude sampling range.

**Fig 5.**
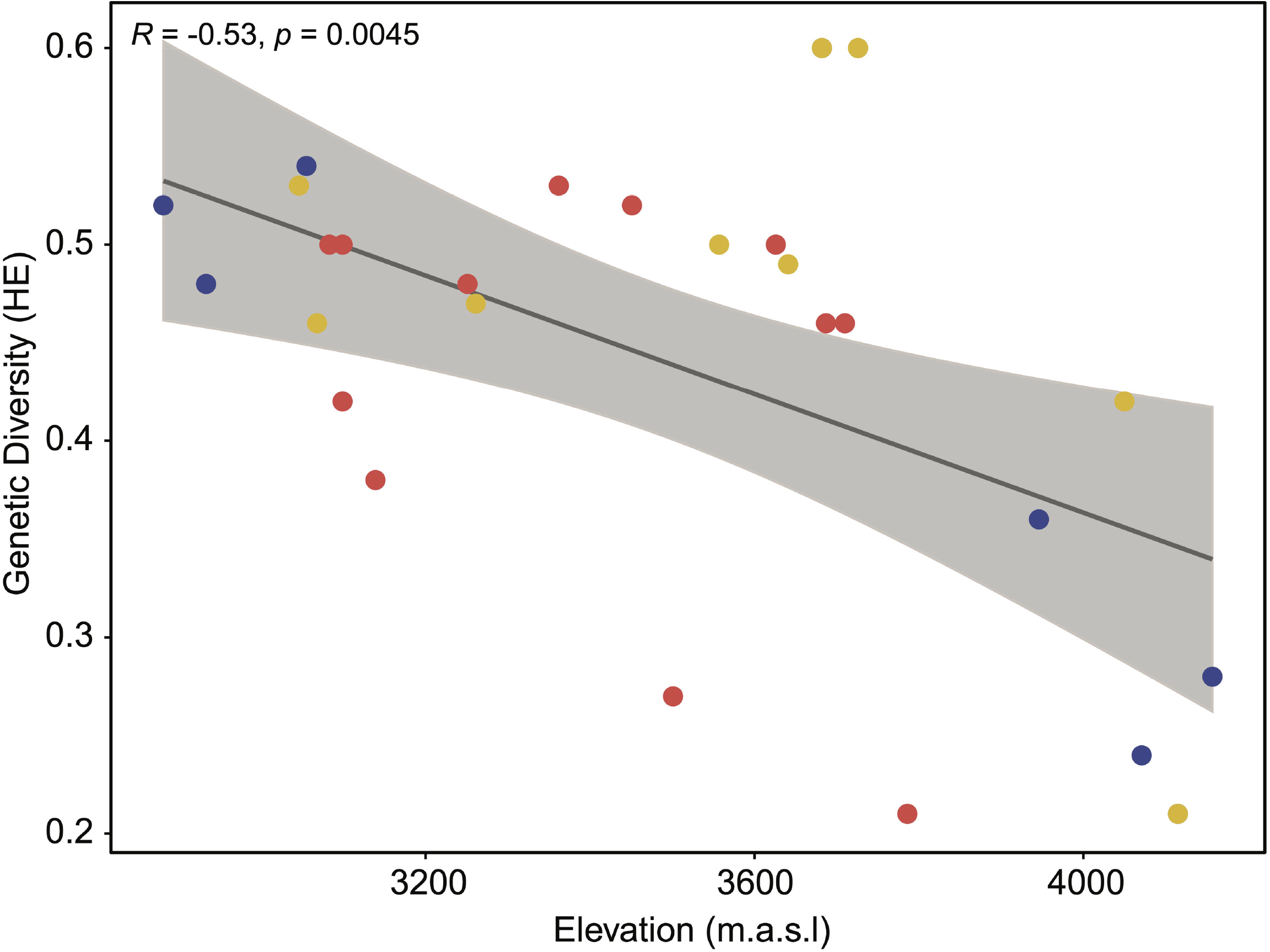
Linear regression between *V. floribundum* genetic diversity and elevation. The plot shows the correlation between elevation (x-axis) and the genetic diversity (y-axis), assessed through the expected heterozygosity estimates per collection sites (confidence interval=95%). Each data point represents one collection site where the individuals were sampled in this study (total of 27 collection sites).

## Discussion

### Genetic diversity of *V. floribundum* in the Ecuadorian Highlands

The mean number of alleles (N_A_=11.2 alleles per locus) and the global expected heterozygosity (H_E_=0.73) found in this study reveal a high degree of genetic diversity for *V. floribundum* in the Ecuadorian Highlands. In a previous study, where we assessed the genetic diversity of *V. floribundum* in the north of Ecuador, we reported on average 6.1 alleles per locus and an expected heterozygosity (H_E_) of 0.49, representing a moderate degree of genetic diversity [36]. Differences in both the number of alleles and the expected heterozygosity between these studies could be explained by the use of heterologous markers (designed for *V. corymbosum*) and the collection of individuals in a limited geographical area. The use of heterologous markers could generate the presence of null alleles that tend to bias the estimation of heterozygosity and inflate F_ST_ values [57–59]. Furthermore, the geographical range and distribution of populations have a significant impact on the amount of genetic variation of a species [60] and could explain the higher degree of genetic diversity found in this study. This can be particularly true for instances where geographically distant individuals are also genetically more distinct, highlighting the importance of sampling range as well as density of collection sites within a geographical region. A greater genetic variability has been previously reported for species distributed over a wide geographical range with respect to those found in a specific limited area [61–63].

The total number of alleles obtained for *V. floribundum* in this study is similar to those reported for wild *V. angustifolium* and *V. corymbosum* clones [35], and higher compared to those reported for *V. macrocarpon* cultivars [64]. Several studies report that intensive selection and inbreeding during domestication processes reduce the genetic diversity in many plant species and increase genetic drift [65,66], which could explain why wild *Vaccinium* species are more diverse than their domesticated counterparts. The global expected heterozygosity (He) in this study is similar to the He reported for *V. sieboldii* (H_E_=0.73) and *V. ciliatum* (H_E_=0.75), both wild endangered species [67]. A high level of genetic diversity in these species could indicate random mating among individuals, a common feature of wild plant species [66–67].

### Population structure and genetic differentiation

The Bayesian approach in STRUCTURE showed that *V. floribundum* individuals were grouped into four genetic clusters (Fig 3). From these, only individuals from cluster 4 also grouped separately in the PCoA (Fig 2), and greatly differentiated from the other clusters based on genetic distances (0.171≤F_ST_≤0.249). This cluster comprises individuals from CS12 (Quilotoa) in the central region, and CS22 (Toreadora), CS23 (Cajas) and CS24 (Cruces) in the southern region. Beyond their genetic similarities, these individuals share the uppermost elevations among the collection sites included in this study (3929-4160 masl) (S1 Table). This higher strata of the *páramo* altitudinal range has been described as a distinct ecosystem known as *superpáramo* [68,69], and its stark ecological differences compared to the lower altitude settings could account for some of the observed differences in the *mortiño* populations established here. First, the effects of elevation on the genetic diversity and differentiation on the species are highlighted by our results that show a correlation between the expected heterozygosity of individuals in specific collection sites and their elevation (*r*=0.29; *p*=0.0001), with (the high-altitude) cluster 4 presenting the lowest expected heterozygosity (H_E_=0.27±0.06) and exclusive allele number (N_PA_=8) (Table 3). Observations of two woody plant species growing in altitudinal clines (*Castanopsis eyeri* and *Daphniphyllum oldhamii*) suggest that factors like temperature, phenological characteristics, low population densities and small effective population sizes could play a role in the isolation and reduction of genetic diversity of populations at higher elevations [70].

Furthermore, the evident differentiation of this cluster of high-elevation populations can be explained by an isolation by elevation model where gene flow from lower to higher elevations is reduced [70,71]. Different mechanisms can drive this altitude-dependent isolation, such as reproductive barriers due to phenological shifts that results in differences in flowering time [69,71]. It is also possible that altitude clines affect pollinator behavior: pollinator visits are less frequent at higher altitudes, which could explain the results obtained in the migration network where the gene flow was minimal and unidirectional from cluster 4 to the other clusters (0.21-0.39) (Fig 4). As a consequence of these types of phenomena, habitats at higher elevations (like in the *superpáramo*) separated by lower elevation valleys can behave like islands where gene flow between different locations is decreased. This kind of “sky island” dynamics can further explain the larger divergence between the populations from cluster 4 and the rest of the *mortiño* genetic diversity in the Ecuadorian Andes.

Nonetheless, the questions of the similarities between geographically distant locations within cluster 4 (i.e. CS12 versus CS22, CS23 and CS24) and the genetic patterns observed in the remaining clusters persist. These patterns might be associated with the geological history and geography of the Andean region, important forces that drive the evolution and distribution of different species in the region [72,73]. In accordance with our previous study [36], the Mantel test results confirmed a significant correlation between genetic and geographic distances (*r*=0.29; *p*=0.0001), suggesting that an isolation-by-geographic distance model also explains part of the population structure of *V. floribundum* in Ecuador. Under this model, physical barriers could play an additional role in the reduced genetic flow between clusters [74,75], particularly between the first three genetic clusters which show moderate to high differentiation (0.078 ≤ F_ST_ ≤ 0.120) (Table 2). The moderate degree of differentiation (F_ST_=0.078) between clusters 1 and 2 could be explained by the emergence of mountainous barriers in the Andean Highlands during the Miocene, as we previously suggested [36] based on results which are consistent with the current study. However, an alternative hypothesis could be related to the spread of ice masses throughout the Andean region during the Pleistocene, a phenomenon that has been proposed as a factor that directly affected the diversity and structure of different species [76,77]. The contraction of the *páramos* during this period could have caused *V. floribundum* populations to be separated by ice masses, reducing or preventing their gene flow and partially isolating both clusters [78]. During the glacial retreat (i.e. melting of the ice masses), individuals from clusters 1 and 2 could have come into contact again, a phenomenon known as secondary contact [77,79].

A third driving force of the observed population structure might include the role of adaptation of *V. floribundum* to different ecosystems, prompting both diversifying selection and convergent evolution. For instance, the physical characteristics of different soil types, determined by their distinct geological origins, partially overlap with the distribution of specific genetic clusters. For instance, individuals from clusters 1 and 2 grow over Andisols developed from volcanic projections characteristic to the northern Ecuadorian highlands, encompassing the northern region and most of the central region in this study (CS1-18). The presence of multiple volcanoes and eolian deposits of volcanic material in these areas have generated a distinct pyroclastic layer over the soils [76]. In contrast, individuals from cluster 3 are found on a wide monotonous plateau. These individuals grow on an old ferritic soil base, also known as metamorphic rock, which is characterized by the accumulation of clay [76]. The geological origin of the soils could then explain some of the differentiation between cluster 3 and both clusters 1 and 2 (F_ST1_=0.114; F_ST2_=0.120), as the unique characteristics between the volcanic and metamorphic rock could act as a force for diversifying selection. Similarly, extreme climatic conditions at higher elevations (characterized by low temperatures and limited nutrient availability [69,80]) can serve as effective selective pressures. Plants adapt to these conditions through strategies such as size reduction, clonal propagation that favors survival over long periods of climatic oscillation, and the fixation of specific alleles [81–83]. This kind of adaptive process could explain the genetic similarities of geographically distant populations within cluster 4 (Fig 2) even in the absence of gene flow. However, the interplay between isolation-by-distance and isolation-by-elevation scenarios is probably more nuanced, as studies have reported that under the latter, gene flow is higher between sites at similar elevations than across elevation clines [71,84].

Future studies should address the occurrence of different adaptations in *mortiño* individuals living in specific habitats (such as the *superpáramo*) and explore the relationship between genetic diversity and the origin of the soils where the *mortiño* grows.

## Conclusions

The genetic data presented in this study demonstrates that *V. floribundum* displays a high degree of genetic diversity in the Ecuadorian Highlands. Population structure analyses revealed that individuals grouped into four genetic clusters, distributed according to their geographic location (northern, central and southern regions). While genetic distances are associated with geographic distances between individuals, it is noteworthy that that elevation also has an effect on the genetic diversity of the populations. Interestingly, the group identified in the PCoA (comprising individuals from CS12, CS22, CS23 and CS24) was consistent with the cluster 4, found through the bayesian analysis. Individuals from this cluster were collected at higher altitudes, presented lower genetic diversity and a higher degree of genetic differentiation from the remaining populations, as well as low gene flow with the populations assigned to the other clusters. It is evident that the structure of *V. floribundum* populations in Ecuador was driven by an interplay between isolation-by-distance and isolation-by-elevation dynamics, and the geography of the Andean region possibly plays a key role in driving these dynamics through a combination of different geographic features, historical climatic fluctuations and the establishment of scenarios favoring the action of adaptive processes.

Furthermore, the high values of genetic diversity that were found are encouraging for the conservation of this species. Since this study assessed the genetic diversity and population structure of wild *V. floribundum* in the ten provinces that encompass the Ecuadorian Highlands, the results presented here could serve as a basis for the development of conservation programs for this species and its habitat at the *páramo*, a relevant and unique ecosystem.

## Acknowledgments

This research was funded with a Chancellor’s Grant from Universidad San Francisco de Quito USFQ (Quito-Ecuador). Genetic data for specimens were obtained under the Genetic Resources Permit Number: MAE-DNB-CM-2016-0046 granted to Universidad San Francisco de Quito by Ministerio del Ambiente Ecuador, in accordance with the Ecuadorian law. We would like to thank Diego Urquía for his contribution in data processing, as well as Dario Ramirez and Andrea Pinos for their assistance in the collection of the samples. We also thank all the members of the Plant Biotechnology Laboratory (Universidad San Francisco de Quito) who contributed to this research.

## Author Contributions

**Conceptualization:** María de Lourdes Torres.

**Data curation:** Pamela Vega-Polo, María Mercedes Cobo.

**Formal analysis:** Pamela Vega-Polo, María Mercedes Cobo, Bernardo Gutierrez.

**Funding acquisition:** María de Lourdes Torres.

**Investigation:** Pamela Vega-Polo, Andrea Argudo, María de Lourdes Torres.

**Methodology:** Pamela Vega-Polo, Andrea Argudo, María Mercedes Cobo, Jennifer Rowntree, María de Lourdes Torres.

**Project administration:** María Mercedes Cobo, María de Lourdes Torres.

**Resources:** Jennifer Rowntree, María de Lourdes Torres.

**Software:** Pamela Vega-Polo, Bernardo Gutierrez.

**Supervision:** María Mercedes Cobo, María de Lourdes Torres.

**Validation:** Bernardo Gutierrez.

**Visualization:** Pamela Vega-Polo, María Mercedes Cobo, Bernardo Gutierrez.

**Writing – original draft:** Pamela Vega-Polo, Andrea Argudo, Jennifer Rowntree.

**Writing – review & editing:** Pamela Vega-Polo, María Mercedes Cobo, Bernardo Gutierrez, María de Lourdes Torres.

## Supporting Information

**S1 Table. Information for the 27 *V. floribundum* collection sites (CS) from 3 defined regions in the Ecuadorian Highlands.**

**S2 Table. List of the 30 SSR homologous markers designed for *V. floribundum*, for this study.**

**S3 Table. Estimate of null allele frequencies for each analyzed *V. floribundum* genetic cluster and SSR locus.** Mean: the mean frequencies over the four genetic clusters.

**S4 Table. Summary of the diversity parameters for each collection site where *V. floribundum* individuals were collected.** N_A_ (number of alleles); N_PA_ (number of private alleles); A_R_ (mean allelic richness calculated using rarefaction); H_O_ (observed heterozygosity); H_E_ (expected heterozygosity); and F_IS_ (fixation index) are given for all 100 individuals genotyped at 16 SSR loci across 27 collection sites in three geographic regions.

**S5 Table. Values for estimating the optimum K from the analysis in STRUCTURE with an admixture model.**

**S6 Table. Results of the analysis of molecular variance (AMOVA) performed for the *V. floribundum* genetic clusters that were identified in the population structure analysis (*P*=0.001).**

**S7 Table. Output for general linear model for the effect of the elevation over the heterozygosity.**

## References

1. Luteyn JL. Diversity, adaptation, and endemism in neotropical ericaceae: Biogeographical patterns in the vaccinieae. Botanical Review. 2002. doi: 10.1663/0006-8101(2002)068[0055:DAAEIN]2.0.CO;2

2. Song GQ, Hancock J. Vaccinium. In: Kole C, editor. Wild Crop Relatives: Genomic and Breeding Resources: Temperate Fruits. Berlin: Springer-Verlag Berlin Heidelberg; 2011. p. 197–221. doi: 10.1007/978-3-642-20447-0

3. Luby JJ, Ballington JR, Draper AD, Pliszka K, Austin ME. BLUEBERRIES AND CRANBERRIES (VACCINIUM). Acta Hortic. 1991. doi: 10.17660/actahortic.1991.290.9

4. Ortiz J, Marín-Arroyo MR, Noriega-Domínguez MJ, Navarro M, Arozarena I. Color, phenolics, and antioxidant activity of blackberry (Rubus glaucus Benth.), blueberry (Vaccinium floribundum Kunth.), and apple wines from ecuador. J Food Sci. 2013. doi: 10.1111/1750-3841.12148

5. Abreu OA, Barreto G, Prieto S. Vaccinium (ericaceae): Ethnobotany and pharmacological potentials. Emirates J Food Agric. 2014;26(7):577–91. doi: 10.9755/ejfa.v26i7.16404

6. Alarcón-Barrera KS, Armijos-Montesinos DS, García-Tenesaca M, Iturralde G, Jaramilo-Vivanco T, Granda-Albuja MG, et al. Wild Andean blackberry (Rubus glaucus Benth) and Andean blueberry (Vaccinium floribundum Kunth) from the Highlands of Ecuador: Nutritional composition and protective effect on human dermal fibroblasts against cytotoxic oxidative damage. J Berry Res. 2018. doi: 10.3233/JBR-180316

7. Lila MA. Impact of Bioflavonoids from Berryfruits on Biomarkers of Metabolic Syndrome. Funct Foods Heal Dis. 2011. doi: 10.31989/ffhd.v1i2.143

8. Llivisaca S, Manzano P, Ruales J, Flores J, Mendoza J, Peralta E, et al. Chemical, antimicrobial, and molecular characterization of mortiño (Vaccinium floribundum Kunth) fruits and leaves. Food Sci Nutr. 2018. doi: 10.1002/fsn3.638

9. Prencipe FP, Bruni R, Guerrini A, Rossi D, Benvenuti S, Pellati F. Metabolite profiling of polyphenols in Vaccinium berries and determination of their chemopreventive properties. J Pharm Biomed Anal. 2014. doi: 10.1016/j.jpba.2013.11.016

10. Schreckinger ME, Wang J, Yousef G, Lila MA, De Mejia EG. Antioxidant capacity and in Vitro inhibition of adipogenesis and inflammation by phenolic extracts of Vaccinium floribundum and Aristotelia chilensis. J Agric Food Chem. 2010. doi: 10.1021/jf100975m

11. Schreckinger M, Lila MA, Yousef G, De Mejia E. Inhibition of α-glucosidase and α-amylase by Vaccinium floribundum and Aristotelia chilensis proanthocyanidins. ACS Symp Ser. 2012. doi: 10.1021/bk-2012-1109.ch006

12. Vasco C, Riihinen K, Ruales J, Kamal-Eldin A. Chemical composition and phenolic compound profile of mortiño (vaccinium floribundum kunth). J Agric Food Chem. 2009. doi: 10.1021/jf9013586

13. Chaparro de Valencia M, Becerra de Lozano N. Anatomia del fruto de Vaccinium floribundum (Ericaceae). Acta Biológica Colomb. 2012;4(1):47–60. doi: 900-1649 0120-548X

14. Magnitskiy S, Ligarreto G, Lancheros H. Rooting of two types of cuttings of fruit crops Vaccinium floribundum Kunth and Disterigma alaternoides (Kunth) Niedenzu (Ericaceae). Agron Colomb. 2011;29:191–203.

15. De la Torre L, Navarrete H, Muriel P, Macía MJ, Balslev H. Enciclopedia de las Plantas Útiles del Ecuador (con extracto de datos). Quito: Herbario QCA de la Escuela de Ciencias Biológicas de la Pontificia Universidad Católica del Ecuador & Herbario AAU del Departamento de Ciencias Biológicas de la Universidad de Aarhus. 2008.

16. Ramsay P, Oxley E. Fire temperatures and postfire plant community dynamics in Ecuadorian grass paramo. Vegetatio. 1996;124(2):129–44. doi: 10.1007/BF00045489

17. Hofstede R, Coppus R, Mena Vasconez P, Segarra P, Sevink J. The conservation status of tussock grass paramo in Ecuador. Ecotropicos. 2002;15(1):3–18.

18. Verweij PA. Spatial and temporal modelling of vegetation pattems-buming and grazing in the paramo of Los Nevados National Park, Colombia. The Netherlands, ITC. 1995.

19. Astudillo PX, Schabo DG, Siddons DC, Farwig N. Patch-matrix movements of birds in the páramo landscape of the southern Andes of Ecuador. Emu - Austral Ornithol. 2019;119(1):53–60. doi: 10.1080/01584197.2018.1512371

20. Echeverry M a., Harper GJ. Fragmentación y deforestación como indicadores del estado de los ecosistemas en el Corredor de Conservación Choco-Manabí (Colombia-Ecuador). Recur Nat y Ambient. 2009;58:78–88.

21. Ellenberg H. Man’s Influence on Tropical Mountain Ecosystems in South America: The Second Tansley Lecture. J Ecol. 1979. doi: 10.2307/2259105

22. Peters T, Drobnik T, Meyer H, Rankl M, Richter M, Rollenbeck R, et al. Environmental Changes Affecting the Andes of Ecuador. In: Ecosystem Services, Biodiversity and Environmental Change in a Tropical Mountain Ecosystem of South Ecuador. 2013. doi: 10.2307/2259105

23. Suárez E, Medina G. Vegetation structure and soil properties in Ecuadorian páramo grasslands with different histories of burning and grazing. Arctic, Antarct Alp Res. 2001;33(2):158–64. doi: 10.1080/15230430.2001.12003418

24. Aguilar R, Quesada M, Ashworth L, Herrerias-Diego Y, Lobo J. Genetic consequences of habitat fragmentation in plant populations: Susceptible signals in plant traits and methodological approaches. Mol Ecol. 2008;17(24):5177–88. doi: 10.1111/j.1365-294X.2008.03971.x

25. Cain ML, Milligan BG, Strand AE. Long-distance seed dispersal in plant populations. Am J Bot. 2000;87(9):1217–27. doi: 10.2307/2656714

26. Couvet D. Deleterious effects of restricted gene flow in fragmented populations. Conserv Biol. 2002;16(2):369–76. doi: 10.1046/j.1523-1739.2002.99518.x

27. Haddad NM, Brudvig LA, Clobert J, Davies KF, Gonzalez A, Holt RD, et al. Habitat fragmentation and its lasting impact on Earth’s ecosystems. Sci Adv. 2015;1(2):e1500052. doi: 10.1126/sciadv.1500052

28. Jacquemyn H, Brys R, Hermy M. Patch occupancy, population size and reproductive success of a forest herb (Primula elatior) in a fragmented landscape. Oecologia. 2002;130(4):617–25. doi: 10.1007/s00442-001-0833-0

29. Young A, Boyle T, Brown T. The population genetic consequences of habitat fragmentation for plants. Trends Ecol Evol. 1996;11(10):413–8. doi: 10.1016/0169-5347(96)10045-8

30. Boches PS, Bassil N V., Rowland LJ. Microsatellite markers for Vaccinium from EST and genomic libraries. Mol Ecol Notes. 2005;5(3):657–60. doi: 10.1111/j.1471-8286.2005.01025.x

31. Debnath SC. An assessment of the genetic diversity within a collection of wild cranberry (Vaccinium macrocarpon Ait.) clones with RAPD-PCR. Genet Resour Crop Evol. 2007;54(3):509–17. doi: 10.1007/s10722-006-0007-3

32. Debnath SC. Development of ISSR markers for genetic diversity studies in Vaccinium angustifolium. Nord J Bot. 2009;27(2):141–8. doi: 10.1111/j.1756-1051.2009.00402.x

33. Gawroński J, Kaczmarska E, Dyduch-Siemińska M. Assessment of genetic diversity between Vaccinium corymbosum L. Cultivars using RAPD and ISSR markers. Acta Sci Pol Hortorum Cultus. 2017;16(3):129–40. doi: 10.24326/asphc.2017.3.13

34. Persson HA, Gustavsson BA. The extent of clonality and genetic diversity in lingonberry (Vaccinium vitis-idaea L.) revealed by RAPDs and leaf-shape analysis. Mol Ecol. 2001;10(6):1385–97. doi: 10.1046/j.1365-294X.2001.01280.x

35. Tailor S, Bykova N V., Igamberdiev AU, Debnath SC. Structural pattern and genetic diversity in blueberry (Vaccinium) clones and cultivars using EST-PCR and microsatellite markers. Genet Resour Crop Evol. 2017;64(8):2071–82. doi: 10.1007/s10722-017-0497-1

36. Cobo MM, Gutiérrez B, Torres AF, Torres M de L. Preliminary analysis of the genetic diversity and population structure of mortiño (Vaccinium floribundum Kunth). Biochem Syst Ecol. 2016;64:14–21. doi: 10.1016/j.bse.2015.11.008

37. Xin Z, Chen J. Extraction of genomic DNA from plant tissues. In: Kieleczawa J, editor. DNA Sequencing II: Optimizing Preparation and Cleanup. Boston: Jones & Bartlett Learning; 2006. p. 47–59.

38. Griffiths SM, Fox G, Briggs PJ, Donaldson IJ, Hood S, Richardson P, et al. A Galaxy-based bioinformatics pipeline for optimised, streamlined microsatellite development from Illumina next-generation sequencing data. Conserv Genet Resour. 2016;8(4):481–6. doi: 10.1007/s12686-016-0570-7

39. Blacket MJ, Robin C, Good RT, Lee SF, Miller AD. Universal primers for fluorescent labelling of PCR fragments-an efficient and cost-effective approach to genotyping by fluorescence. Mol Ecol Resour. 2012;12(3):456–63. doi: 10.1111/j.1755-0998.2011.03104.x

40. Clark L V., Jasieniuk M. polysat: An R package for polyploid microsatellite analysis. Mol Ecol Resour. 2011;11(3):562–6. doi: 10.1111/j.1755-0998.2011.02985.x

41. Jombart T. Adegenet: A R package for the multivariate analysis of genetic markers. Bioinformatics. 2008;24(11):1403–5. doi: 10.1093/bioinformatics/btn129

42. Goudet J. HIERFSTAT, a package for R to compute and test hierarchical F-statistics. Mol Ecol Notes. 2005;5(1):184–6. doi: 10.1111/j.1471-8286.2004.00828.x

43. Kamvar ZN, Tabima JF, Gr□unwald NJ. Poppr: An R package for genetic analysis of populations with clonal, partially clonal, and/or sexual reproduction. PeerJ. 2014;2:e281. doi: 10.7717/peerj.281

44. Chapuis MP, Estoup A. Microsatellite null alleles and estimation of population differentiation. Mol Biol Evol. 2007. doi: 10.1093/molbev/msl191

45. Keenan K, Mcginnity P, Cross TF, Crozier WW, Prodöhl PA. DiveRsity: An R package for the estimation and exploration of population genetics parameters and their associated errors. Methods Ecol Evol. 2013;4(8):782–8. doi: 10.1111/2041-210X.12067

46. Dray S, Dufour AB. The ade4 package: Implementing the duality diagram for ecologists. J Stat Softw. 2007;22(4):1–20. doi: 10.18637/jss.v022.i04

47. Beck MW. ggord: Ordination Plots with ggplot2. R package version 1.0.0. [Internet]. 2017. Available from: https://zenodo.org/badge/latestdoi/35334615

48. Pritchard JK, Stephens M, Donnelly P. Inference of population structure using multilocus genotype data. Genetics. 2000;155(2):945–59.

49. Evanno G, Regnaut S, Goudet J. Detecting the number of clusters of individuals using the software STRUCTURE: A simulation study. Mol Ecol. 2005;14(8):2611–20. doi: 10.1111/j.1365-294X.2005.02553.x

50. Earl DA, VonHoldt BM. STRUCTURE HARVESTER: A website and program for visualizing STRUCTURE output and implementing the Evanno method. Conserv Genet Resour. 2012;4(2):359–61. doi: 10.1007/s12686-011-9548-7

51. Jakobsson M, Rosenberg NA. CLUMPP: A cluster matching and permutation program for dealing with label switching and multimodality in analysis of population structure. Bioinformatics. 2007;3(14):1801–6. doi: 10.1093/bioinformatics/btm233

52. Rosenberg NA. DISTRUCT: A program for the graphical display of population structure. Mol Ecol Notes. 2004;4(1):137–8. doi: 10.1046/j.1471-8286.2003.00566.x

53. Weir BS, Cockerham CC. Estimating F-statistics for the analysis of population structure. Evolution (N Y). 1984;1358–70. doi: 10.1111/j.1558-5646.1984.tb05657.x

54. Paradis E, Claude J, Strimmer K. APE: Analyses of phylogenetics and evolution in R language. Bioinformatics. 2004;20(2):289–90. doi: 10.1093/bioinformatics/btg412

55. R Development Core Team R. R: A Language and Environment for Statistical Computing. R Foundation for Statistical Computing. 2013. doi: 10.1007/978-3-540-74686-7

56. Kassambara A. ggpubr: “ggplot2” Based Publication Ready Plots. R package version 0.1.7. [Internet]. 2018. Available from: https://cran.r-project.org/package=ggpubr

57. Guidugli MC, Accoroni KAG, Mestriner MA, Contel EPB, Martinez CA, Alzate-Marin AL. Genetic characterization of 12 heterologous microsatellite markers for the giant tropical tree Cariniana legalis (Lecythidaceae). Genet Mol Biol. 2010;33(1):131–4. doi: 10.1590/S1415-47572010000100022

58. Nazareno AG, Pereira RAS, Feres JM, Mestriner MA, Alzate-Marin AL. Transferability and characterization of microsatellite markers in two Neotropical Ficus species. Genet Mol Biol. 2009;32(3):568–71. doi: 10.1590/S1415-47572009005000056

59. Wang C, Schroeder KB, Rosenberg NA. A maximum-likelihood method to correct for allelic dropout in microsatellite data with no replicate genotypes. Genetics. 2012;192(2):651–69. doi: 10.1534/genetics.112.139519

60. Levy E, Byrne M, Coates DJ, Macdonald BM, McArthur S, Van Leeuwen S. Contrasting influences of geographic range and distribution of populations on patterns of genetic diversity in two sympatric Pilbara acacias. PLoS One. 2016;11(10):e0163995. doi: 10.1371/journal.pone.0163995

61. Gitzendanner MA, Soltis PS. Patterns of genetic variation in rare and widespread plant congeners. Am J Bot. 2000;87(6):783–92. doi: 10.2307/2656886

62. Hamrick JL, Godt MJW, Sherman-Broyles SL. Factors influencing levels of genetic diversity in woody plant species. Population Genetics of Forest Trees. 1992. doi: 10.1007/BF00120641

63. Karron JD. A comparison of levels of genetic polymorphism and self-compatibility in geographically restricted and widespread plant congeners. Evol Ecol. 1987;1(1):47–58. doi: 10.1007/BF02067268

64. Fajardo D, Morales J, Zhu H, Steffan S, Harbut R., Bassil N, et al. Discrimination of American Cranberry Cultivars and Assessment of Clonal Heterogeneity Using Microsatellite Markers. Plant Mol Biol Report. 2013;31(2):264–71. doi: 10.1007/s11105-012-0497-4

65. Doebley JF, Gaut BS, Smith BD. The Molecular Genetics of Crop Domestication. Cell. 2006;127:1309–21. doi: 10.1016/j.cell.2006.12.006

66. Gao Y, Yin S, Wu L, Dai D, Wang H, Liu C, et al. Genetic diversity and structure of wild and cultivated Amorphophallus paeoniifolius populations in southwestern China as revealed by RAD-seq. Sci Rep. 2017;7(1):1–10. doi: 10.1038/s41598-017-14738-6

67. Hirai M, Yoshimura S, Ohsako T, Kubo N. Genetic diversity and phylogenetic relationships of the endangered species Vaccinium sieboldii and Vaccinium ciliatum (Ericaceae). Plant Syst Evol. 2010;287(1-2):75–84. doi: 10.1007/s00606-010-0291-4

68. Salgado-Labouriau ML, Rull V, Schubert C, Va;astro S. The establishment of vegetation after late pleistocene deglaciation in the Paramo de Miranda, Venezuelan Andes. Rev Palaeobot Palynol. 1988;55(1-3):5–17. doi: 10.1016/0034-6667(88)90052-8

69. Sklenář P, Balslev H. Superpáramo plant species diversity and phytogeography in Ecuador. Flora Morphol Distrib Funct Ecol Plants. 2005;5:416–33. doi: 10.1016/j.flora.2004.12.006

70. Hahn CZ, Michalski SG, Fischer M, Durka W. Genetic diversity and differentiation follow secondary succession in a multi-species study on woody plants from subtropical China. J Plant Ecol. 2017;10(1):213–21. doi: 10.1093/jpe/rtw054

71. Shi MM, Michalski SG, Chen XY, Durka W. Isolation by elevation: Genetic structure at neutral and putatively non-neutral loci in a dominant tree of subtropical forests, castanopsis eyrei. PLoS One. 2011;6(6):e21302. doi: 10.1371/journal.pone.0021302

72. Hazzi NA, Moreno JS, Ortiz-Movliav C, Palacio RD. Biogeographic regions and events of isolation and diversification of the endemic biota of the tropical Andes. Proc Natl Acad Sci U S A. 2018;115(31):7985–90. doi: 10.1073/pnas.1803908115

73. Schmid R., Luteyn JL. Páramos: A Checklist of Plant Diversity, Geographical Distribution, and Botanical LiteratureParamos: A Checklist of Plant Diversity, Geographical Distribution, and Botanical Literature. Taxon. 1999. doi: 10.2307/1224592

74. Bell DJ, Rowland LJ, Zhang D, Drummond FA. Spatial genetic structure of lowbush blueberry, Vaccinium angustifolium, in four fields in Maine. Botany. 2009;87(10):932–46. doi: 10.1139/B09-058

75. Diniz-Filho JAF, Soares TN, Lima JS, Dobrovolski R., Landeiro VL, Telles MP de C, et al. Mantel test in population genetics. Genet Mol Biol. 2013;36(4):475–85. doi: 10.1590/S1415-47572013000400002

76. Moreno J, Yerovi F, Herrera M, Yanez D, Espinosa J. Soils from the Highlands. In: Espinosa J, Moreno J, Bernal G, editors. The Soils of Ecuador. Springer. World Soils Book Series, Springer Cham; 2016. p. 79–137. doi: 10.1007/978-3-319-25319-0_3

77. Zemlak TS, Habit EM, Walde SJ, Battini MA, Adams EDM, Ruzzante DE. Across the southern Andes on fin: Glacial refugia, drainage reversals and a secondary contact zone revealed by the phylogeographical signal of Galaxias platei in Patagonia. Mol Ecol. 2008;17(23):5049–61. doi: 10.1111/j.1365-294X.2008.03987.x

78. Vuilleumier BS. Pleistocene Changes in the fauna and flora of South America. Science (80-). 1971;173(4):771–80. doi: 10.1126/science.173.3999.771

79. Nevado B, Contreras-Ortiz N, Hughes C, Filatov DA. Pleistocene glacial cycles drive isolation, gene flow and speciation in the high-elevation Andes. New Phytol. 2018;219(2):779–93. doi: 10.1111/nph.15243

80. Luteyn JL. Paramo: an Andean ecosystem under human influence. 1^st^ ed. London: UK Academic Press; 1992.

81. Linhart YB, Gehring JL. Genetic Variability and Its Ecological Implications in the Clonal Plant Carex scopulorum Holm. in Colorado Tundra. Arctic, Antarct Alp Res. 2003;35(4):429–33. doi: 10.1657/1523-0430(2003)035[0429:GVAIEI]2.0.CO;2

82. Peng Y, Macek P, Macková J, Romoleroux K, Hensen I. Clonal Diversity and Fine-scale Genetic Structure in a High Andean Treeline Population. Biotropica. 2015;47(1):59–65. doi: 10.1111/btp.12175

83. Wesche K, Hensen I, Undrakh R. Genetic structure of Galitzkya macrocarpa and G. potaninii, two closely related endemics of central Asian mountain ranges. Ann Bot. 2006;98(5):1025–34. doi: 10.1093/aob/mcl182

84. Byars SG, Parsons Y, Hoffmann AA. Effect of altitude on the genetic structure of an Alpine grass, Poa hiemata. Ann Bot. 2009;103(6):885–99. doi: 10.1093/aob/mcp018

