## Supplementary material for "Characterizing the genetic diversity of the Andean blueberry (*Vaccinium floribundum* Kunth.) across the Ecuadorian Highlands": S1 Table

**S1 Table. Information for the 27 *V. floribundum* collection sites (CS) from 3 defined regions in the Ecuadorian Highlands.**

| Region | Collection site | Location | Province | Number of individuals | Elevation |
|---|---|---|---|---|---|
| Northern | CS1 | La Cofradia | Carchi | 3 | 3243-3260 |
| | CS2 | San Gabriel | Carchi | 3 | 3405-3489 |
| | CS3 | El Angel | Carchi | 4 | 3317-3402 |
| | CS4 | Cahuasquí | Imbabura | 4 | 3588-3714 |
| | CS5 | Cuicocha | Imbabura | 4 | 3080-3122 |
| | CS6 | Santa Lucía | Imbabura | 3 | 2994-3062 |
| | CS7 | Mojanda | Pichincha | 4 | 3596-3958 |
| | CS8 | Cayambe | Pichincha | 3 | 3721-3885 |
| | CS9 | Lloa | Pichincha | 4 | 3450-3546 |
| | CS10 | PNC | Cotopaxi | 4 | 3550-3803 |
| | CS11 | Sigchos | Cotopaxi | 3 | 3118-3147 |
| Central | CS12 | Quilotoa | Cotopaxi | 4 | 4090-4131 |
| | CS13 | Tisandeo | Tungurahua | 5 | 3542-3601 |
| | CS14 | Carihuairazo | Tungurahua | 5 | 3627-3787 |
| | CS15 | Salinas Norte | Bolivar | 3 | 4038-4062 |
| | CS16 | Salinas | Bolivar | 3 | 3609-3670 |
| | CS17 | Cebapamba | Bolivar | 4 | 3243-3285 |
| | CS18 | Quimiac | Chimborazo | 7 | 3482-3758 |
| | CS19 | Cerro Abuga | Cañar | 3 | 3029-3120 |
| | CS20 | Surimpalti | Cañar | 4 | 3015-3066 |
| Southern | CS21 | San Miguel | Cañar | 3 | 3044-3100 |
| | CS22 | Toreadora | Azuay | 4 | 3929-3956 |
| | CS23 | Cajas | Azuay | 3 | 4057-4098 |
| | CS24 | Cruces | Azuay | 3 | 4154-4160 |
| | CS25 | Saraguro | Loja | 3 | 2881-3008 |
| | CS26 | Santiago | Loja | 3 | 2868-2911 |
| | CS27 | Podocarpus | Loja | 4 | 3039-3067 |
