## Supplementary material for "Characterizing the genetic diversity of the Andean blueberry (*Vaccinium floribundum* Kunth.) across the Ecuadorian Highlands": S2 Table

**S2 Table. List of the 30 SSR homologous markers designed for *V. floribundum*, for this study.**

| Locus | Primer sequences (5'-3') | Motive | T °C |
|---|---|---|---|
| **Mo001** | **F:AACCTGTACAAGTCTACCCCTACCG**<br>**R: TAATAACAGAACATCAGTGCAAGGC** | **(TTCCTG)48** | **58** |
| **Mo002** | **F: CAAAATAACCCTCAAACACACACC**<br>**R: TTTATCATTATCCTACAGCGTCACC** | **(TTTGG)50** | **58** |
| Mo003 | F: CTACCCATCCAACCACCTAACC<br>R: TAATAACAGAACATCAGTGCAAGGC | (ATGG)36 | 58 |
| **Mo004** | **F: GAGGTATTGGAATCCTTGGATGG**<br>**R: CTCTTCCCCTCAACTCTTCCC** | **(TCC)33** | **58** |
| **Mo005** | **F: TAGAGATTCATCTCCATCCTTTTGC**<br>**R: GTCCATTAGGGTTCCAAAAGTGC** | **(ACC)33** | **58** |
| Mo006 | F: GTTTGGAAGATCTCGCTGAGTAGG<br>R: CAGTAGAACCACTTACCCCTGTAGC | (ACC)27 | 63 |
| **Mo007** | **F: GAAGCCTGGTCAGTCCTTTCC**<br>**R: CACTAGGAGTCTGACTTTCCTCTGC** | **(TGC)24** | **63** |
| **Mo008** | **F: ACTACCCTGCCACTCTCACTACC**<br>**R: CGGACCCAGAGTTAGGATAATACC** | **(ACC)27** | **63** |
| **Mo009** | **F: TATTCTTATGTTCGTCCTCGTAGGC**<br>**R: TTTCCTGCTAGCTGTTGTTGTAACG** | **(AGT)24** | **63** |
| **Mo010** | **F: TAGACAACCACTTTCTTTGGTTTCC**<br>**R: AATAAGTCTTGCTTTGTACCTTGCC** | **(TTC)30** | **63** |
| **Mo011** | **F: GCGAGAGTATTGGTGTTCATGC**<br>**R: CAGGTATAGATATACTGGGTTTTGAGG** | **(TTC)24** | **58** |
| Mo012 | F: TGTTACGCTTATTACGTTGTGTTGG<br>R: TTCACAGTTGACTTGTTCTTATGCC | (ACC)24 | 60 |
| Mo013 | F: AGAGTACCATTTGGGTTAGTTTTGC<br>R: GACCAAAACAGTAGAAAACGACAGC | (TTC)33 | 60 |
| Mo014 | F: CTTTAAATGGAACCCCTCTTGTAGG<br>R: AATACATACAATCTCAGGCAAAGGG | (ATT)30 | 60 |
| **Mo015** | **F: TAAATCCAAAAGGACAACTCCATCC**<br>**R: AACATGGGTTTAGCGTAGGAGACG** | **(TTC)27** | **60** |
| **Mo016** | **F: GAAGAAGAAATGGTGAGACAACTGC**<br>**R: AAGAAGATTGACTAGGGAGACATCG** | **(TTC)33** | **60** |
| Mo017 | F: AAAACGTAGTTGGACAAACGATACG<br>R: GGTGGTTGTGGCCAAAATAGG | (ATT)42 | 60 |
| **Mo018** | **F:ATTCGGGTATGGAGAGAGAAAGAGG**<br>**R: ACACCAACAAACCCGAAAATAACC** | **(TCC)27** | **60** |
| Mo019 | F: AACCTGTGTAATCTCACCCACTACC<br>R: ATAGATGAGGTGCAACAAGAGTTGG | (AAC)33 | 58 |
| **Mo020** | **F: CTACATTTTACCCGGTCACTTTTGC**<br>**R: CACACTAGTTACAAGAGCATTTTCCC** | **(ATT)33** | **60** |
| **Mo021** | **F: CATGGTTTGGTCTAGTTGATAACCC**<br>**R: GATGCTTCCTAGAGCCTTTATTGC** | **(TTC)33** | **60** |
| Mo022 | F: CTTTAGAAACACGAAGTGACAGACC<br>R: TAGCTAATAGGTCAAGGGTTCAAGG | (TTC)30 | 60 |
| Mo023 | F: GACGACCAGAATAAAGAAGAGAAAGG<br>R: CTGAAGCCGACATATAGAAACTTGG | (ATT)24 | 60 |
| **Mo024** | **F: TGTTGCTTCTTTTGTTTCCTACCC**<br>**R: TTAGAGTTCTAAGCCAACAAACTCG** | **(ATC)24** | **60** |
| **Mo025** | **F: GGTCAAAAGGAGGAGAATAATAGCC**<br>**R: TGTCTCTGCCCATTTTAATGTTACC** | **(TTC)54** | **60** |
| Mo026 | F: AAGGGGCATCACTGTAATAGTATCG<br>R: GAACGAAATCATCGTCTTCACG | (TC)30 | 60 |
| Mo027 | F: CTTACCATGGTTGTAGCATTTTCCC<br>R: GTCCGTCAATTGTTTATGGATAAGG | (AT)20 | 60 |
| Mo028 | F: TGCTTTCATCTATTGACCTTATCGG<br>R: GGGTGTCGTATCAGTAATCTCTCG | (AC)18 | 60 |
| Mo029 | F: ACACCCTCAACTCAATACAGTGTGC<br>R: TGATGAGAATCCACCATTTTAGTGC | (TC)30 | 60 |
| Mo030 | F: TAGACAACCAAGCTCATTTTGATCG<br>R: CACTTAGAACAGGAACTCACCTTGC | (TC)18 | 60 |
