## Supplementary material for "Characterizing the genetic diversity of the Andean blueberry (*Vaccinium floribundum* Kunth.) across the Ecuadorian Highlands": S3 Table

**S3 Table. Estimate of null allele frequencies for each analyzed *V. floribundum* genetic cluster and SSR locus.**

| Locus | Cluster 1 | Cluster 2 | Cluster 3 | Cluster 4 | Mean |
|---|---|---|---|---|---|
| **Mo001** | 0.089 | 0.131 | 0.088 | 0.091 | 0.100 |
| **Mo002** | 0.000 | 0.039 | 0.016 | 0.081 | 0.034 |
| **Mo004** | 0.275 | 0.215 | 0.017 | 0.117 | 0.156 |
| **Mo005** | 0.320 | 0.194 | 0.086 | 0.068 | 0.167 |
| **Mo007** | 0.224 | 0.142 | 0.052 | 0.048 | 0.117 |
| **Mo008** | 0.001 | 0.065 | 0.113 | 0.129 | 0.077 |
| **Mo009** | 0.352 | 0.174 | 0.215 | 0.106 | 0.211 |
| **Mo010** | 0.236 | 0.139 | 0.129 | 0.121 | 0.156 |
| **Mo011** | 0.001 | 0.126 | 0.048 | 0.000 | 0.044 |
| **Mo015** | 0.001 | 0.216 | 0.077 | 0.297 | 0.148 |
| **Mo016** | 0.342 | 0.160 | 0.171 | 0.000 | 0.168 |
| **Mo018** | 0.261 | 0.145 | 0.198 | 0.174 | 0.195 |
| **Mo020** | 0.176 | 0.150 | 0.104 | 0.224 | 0.163 |
| **Mo021** | 0.369 | 0.120 | 0.162 | 0.069 | 0.180 |
| **Mo024** | 0.330 | 0.089 | 0.091 | 0.082 | 0.148 |
| **Mo025** | 0.453 | 0.178 | 0.214 | 0.055 | 0.225 |

Mean: the mean frequencies over the four genetic clusters.
