## Supplementary material for "Characterizing the genetic diversity of the Andean blueberry (*Vaccinium floribundum* Kunth.) across the Ecuadorian Highlands": S4 Table

**S4 Table. Summary of the diversity parameters for each collection site where *V. floribundum* individuals were collected.**

| Collection site | $N_A$ | $N_{PA}$ | $A_R$ | $H_O$ | $H_E$ | $F_{IS}$ |
|---|---|---|---|---|---|---|
| **CS1** | 43 | - | 2.31 | 0.54 | 0.48 | 0.03 |
| **CS2** | 45 | 2 | 2.28 | 0.54 | 0.52 | 0.12 |
| **CS3** | 49 | - | 2.25 | 0.53 | 0.53 | 0.12 |
| **CS4** | 40 | 4 | 2.04 | 0.47 | 0.5 | 0.07 |
| **CS5** | 40 | 1 | 2.02 | 0.55 | 0.5 | -0.05 |
| **CS6** | 36 | 1 | 1.83 | 0.46 | 0.42 | 0.11 |
| **CS7** | 42 | 3 | 1.97 | 0.34 | 0.46 | 0.33 |
| **CS8** | 28 | 2 | 1.31 | 0.14 | 0.21 | 0.4 |
| **CS9** | 29 | 1 | 1.37 | 0.23 | 0.27 | 0.19 |
| **CS10** | 42 | 2 | 2.05 | 0.42 | 0.46 | 0.15 |
| **CS11** | 37 | - | 1.78 | 0.35 | 0.38 | 0.14 |
| **CS12** | 26 | 1 | 1.23 | 0.07 | 0.21 | 0.78 |
| **CS13** | 53 | 5 | 2.15 | 0.46 | 0.5 | 0.2 |
| **CS14** | 60 | 6 | 2.54 | 0.58 | 0.6 | 0.17 |
| **CS15** | 42 | 1 | 2.16 | 0.46 | 0.42 | 0.08 |
| **CS16** | 42 | 1 | 2.23 | 0.58 | 0.49 | -0.07 |
| **CS17** | 45 | - | 2.1 | 0.5 | 0.47 | -0.01 |
| **CS18** | 65 | 5 | 2.49 | 0.55 | 0.6 | 0.1 |
| **CS19** | 39 | 1 | 1.99 | 0.52 | 0.46 | -0.04 |
| **CS20** | 51 | - | 2.33 | 0.36 | 0.53 | 0.44 |
| **CS21** | 45 | 1 | 2.33 | 0.48 | 0.5 | 0.19 |
| **CS22** | 34 | 2 | 1.65 | 0.09 | 0.36 | 0.82 |
| **CS23** | 26 | 1 | 1.42 | 0.1 | 0.24 | 0.58 |
| **CS24** | 28 | 3 | 1.5 | 0.06 | 0.28 | 0.76 |
| **CS25** | 41 | 1 | 2.03 | 0.52 | 0.48 | 0.06 |
| **CS26** | 48 | 5 | 2.49 | 0.56 | 0.52 | 0.09 |
| **CS27** | 51 | 4 | 2.25 | 0.61 | 0.54 | -0.02 |

$N_A$ (number of alleles); $N_{PA}$ (number of private alleles); $A_R$ (mean allelic richness calculated using rarefaction); $H_O$ (observed heterozygosity); $H_E$ (expected heterozygosity); and $F_{IS}$ (fixation index) are given for all 100 individuals genotyped at 16 SSR loci across 27 collection sites in three geographic regions.
