## Supplementary material for "Characterizing the genetic diversity of the Andean blueberry (*Vaccinium floribundum* Kunth.) across the Ecuadorian Highlands": S5 Table

**S5 Table. Values for estimating the optimum K from the analysis in STRUCTURE with an admixture model.**

| K | Reps | Mean LnP(K) | Stdev LnP(K) | Ln'(K) | \|Ln''(K)\| | Delta K |
|---|---|---|---|---|---|---|
| **3** | 10 | -4796.425 | 0.9500 | 283.750000 | 56.500000 | 59.473684 |
| **4** | 10 | -4569.1750 | 1.0595 | 227.250000 | 117.475000 | 110.879760 |
| **5** | 10 | -4459.4000 | 2.0510 | 109.775000 | 40.025000 | 19.514718 |
