## Supplementary material for "Characterizing the genetic diversity of the Andean blueberry (*Vaccinium floribundum* Kunth.) across the Ecuadorian Highlands": S6 Table

**S6 Table. Results of the analysis of molecular variance (AMOVA) performed for the *V. floribundum* genetic clusters that were identified in the population structure analysis (*P*=0.001).**

| Source | Df | Sum Sq | Mean Sq | Est. Var. | % |
|---|---|---|---|---|---|
| Between clusters | 3 | 159.61 | 53.20 | 1.92 | 21.36% |
| Within clusters | 96 | 677.47 | 7.06 | 7.06 | 78.64% |
| **Total** | 99 | 837.08 | | 8.98 | 100% |
