## Supplementary material for "Characterizing the genetic diversity of the Andean blueberry (*Vaccinium floribundum* Kunth.) across the Ecuadorian Highlands": S7 Table

**S7 Table. Output for general linear model for the effect of the elevation over the heterozygosity.**

| Source | DF | Adj SS | Adj MS | F-Value | P-Value |
|---|---|---|---|---|---|
| Elevation | 1 | 0.090684 | 0.090684 | 9.74 | 0.005 |
| Error | 25 | 0.232723 | 0.009309 | | |
| Lack-of-Fit | 24 | 0.229523 | 0.009563 | 2.99 | 0.432 |
| Pure Error | 1 | 0.003200 | 0.003200 | | |
| Total | 26 | 0.323407 | | | |
